# Optical Limits in Skin Reflectance Measurement: Quantifying Melanin-Dependent Constraints on Erythema Detection

**DOI:** 10.64898/2025.12.22.696093

**Authors:** Yemko Pryor, Junhui He, Jian Kang, Blair Jenkins, Tina Lasisi

## Abstract

As skin color measurement shifts from subjective classification to quantitative assessment, the assumption that objective measures are unbiased requires scrutiny. We provide a physics-informed framework for interpreting visible-range skin measurements, clarifying terminology, describing light-chromophore interactions, and surveying tools and output metrics. The physics of light-tissue interaction constrains what any visible-range observation can reveal: melanin and hemoglobin absorb across overlapping wavelengths, and as melanin increases, hemoglobin’s signature is masked. By analyzing over 15,000 spectra from the International Skin Spectra Archive, we demonstrate that this attenuation is systematic and most severe in the darkest skin, where data are scarcest. These limits arise from the physical properties of skin’s biological constituents and consequently apply to any sensor operating in the visible range. We conclude with guidance on analytical methods, sampling strategies, and extended wavelengths to improve measurement validity across the full range of human skin pigmentation.

**Graphical Abstract:** Objective skin measurements are not inherently unbiased: melanin’s broadband absorption progressively masks hemoglobin’s spectral signature, reducing the sensitivity of visible-range erythema detection in darker skin.

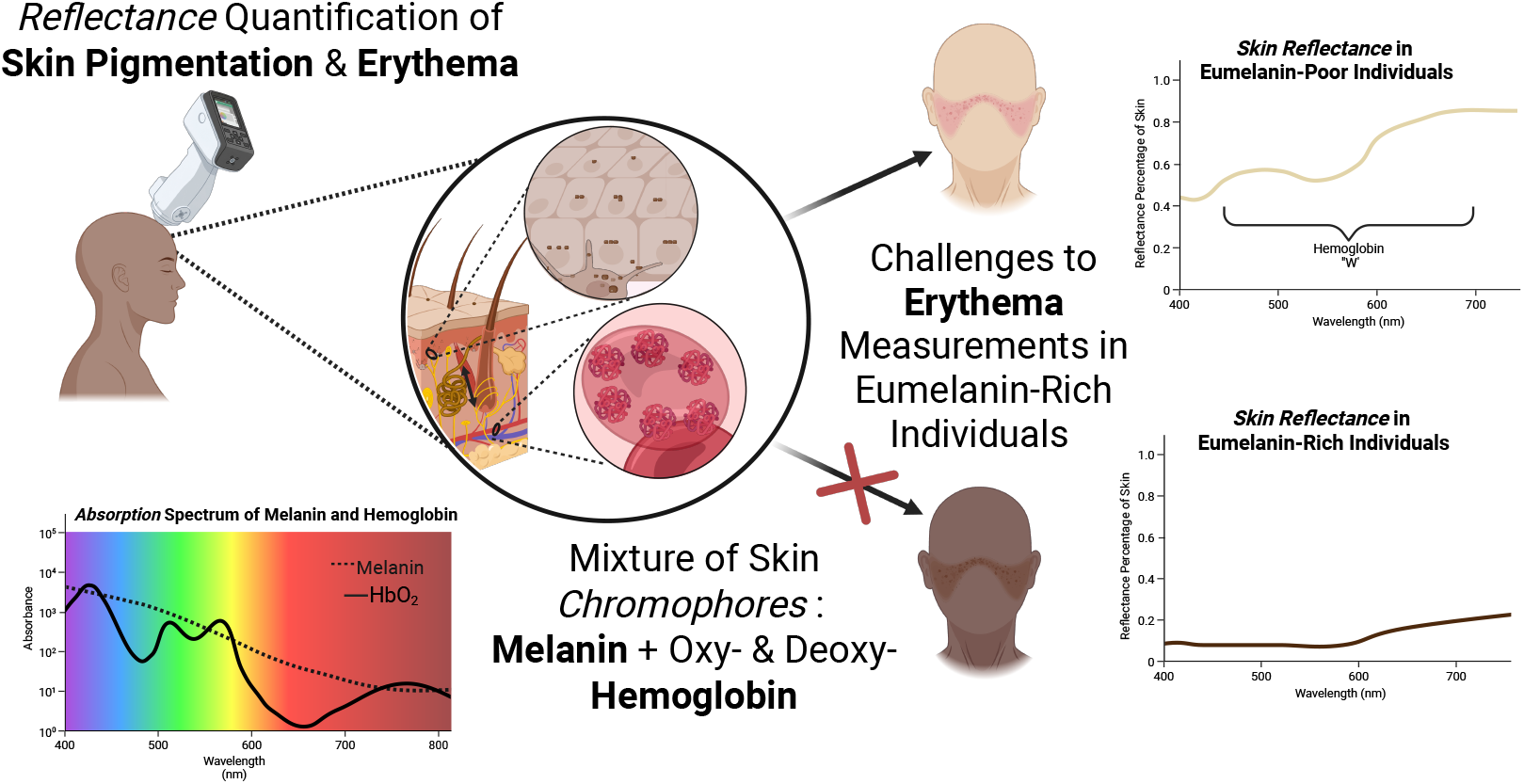

**Summary Points:**

- Visible-range skin reflectance measurements capture real optical information but do not directly measure chromophore concentrations; interpreting these signals requires understanding light-tissue interactions.
- Melanin and hemoglobin absorb at overlapping wavelengths, creating optical entanglement that limits the separability of pigmentation and erythema signals from reflectance data.
- Hemoglobin spectral signatures are systematically attenuated as melanin increases, reducing the sensitivity of erythema detection in darker skin. This represents objective bias arising from physics, not subjective bias in assessment.
- Derivative analysis of full reflectance spectra can reveal hemoglobin-related structure that persists in darker skin even when standard indices show no signal.
- Melanin index distributions should be reported to assess sample representativeness, as demographic diversity does not guarantee adequate coverage of the pigmentation range relevant to optical measurement.

**Limitations:**

- Spectral overlap between melanin and hemoglobin is a physical property of these chromophores; no visible-range instrument or algorithm can fully overcome this constraint.
- These optical limits apply to any visible-range sensor, including spectrophotometers, clinical photography, and AI systems trained on photographic data.
- Existing datasets remain sparse at the high end of the melanin index distribution, limiting validation of measurement approaches in the darkest skin.

## 1 Introduction

Clinical evaluation is beginning to incorporate quantitative assessments of pigmentation and erythema, and researchers are increasingly turning to AI and computer vision tools to extract color-based information from skin. This shift reflects a growing recognition that subjective descriptors, whether visual assessment, self-reported color, or categorical systems such as the Fitzpatrick phototype, cannot capture the continuous variation in pigmentation and vascular responses that underlies skin appearance (Vasudevan et al., 2024; Ly et al., 2020). Objective, instrument-based measurements promise consistency, reproducibility, and the possibility of linking visible phenotype to biological mechanisms. To evaluate what these tools can quantify, however, we must first understand what they are actually measuring.

The transition from subjective to objective approaches does not guarantee that measurements are unbiased or biologically interpretable. Visible-range devices and imaging systems generate numerical values that are often treated as direct representations of underlying tissue properties. In practice, these values reflect how light interacts with skin in situ rather than any single chromophore or physiological parameter. The distinction between perceived skin color and underlying skin pigmentation is therefore essential. Visible appearance is shaped by melanin, hemoglobin, and tissue scattering, determining which photons return to a sensor and thus limit inference from visible-range measurements alone.

For clinicians and researchers, this creates both an opportunity and a challenge. Objective measurements can inform assessments of treatment response, clarify risk associated with pigmentation-related conditions, and enrich research on skin biology and disease prevalence. Without accounting for how skin structure and pigmentation constrain optical measurements, device readouts are easily overinterpreted, leading to systematic bias.

The purpose of this article is to provide a practical, physics-informed framework for interpreting visible-range measurements of skin. We clarify terminology (Table 1), outline how light interacts with skin chromophores, and review commonly used measurement tools. We then present an analysis of multi-population reflectance data from the International Skin Spectra Archive that illustrates how melanin and anatomical site jointly constrain what visible-range measurements can capture, and conclude with guidance for improving measurement validity across the full range of human skin pigmentation.

**Table 1:** Key terminology for skin color measurement.

| Definition | Citations |
| --- | --- |
| **Optics** is the study of how light interacts with skin through absorption, scattering, reflection, and transmission. Optical measurements infer tissue properties from the light remitted from or transmitted through the skin. | (Tuchin, 2015; Van Gemert et al., 1989) |
| **Spectroscopy, spectrophotometry, and spectrometry** are related terms often used interchangeably in practice. Formally, *spectroscopy* refers to the study of light-matter interaction as a function of wavelength, *spectrometry* refers to the measurement process itself, and *spectrophotometry* specifically measures light intensity in controlled reflectance or transmittance geometries. In skin research, terms like "reflectance spectroscopy" and "spectrophotometry" are both used to describe diffuse reflectance measurements. | (Goenaga Infante et al., 2021) |
| **Colorimetry**, sometimes called tristimulus colorimetry, is a color-science method that represents perceived color using three broad spectral channels and standard observer functions, typically expressed as CIE XYZ or CIE L*a*b*. | (Weatherall and Coombs, 1992; Fullerton et al., 1996; Ly et al., 2020) |
| **Melanometry** is the use of optical measurements to quantify melanin-related skin pigmentation, including melanin indices, colorimetric metrics, or spectral fitting. The term "melanometry" was coined recently and is not uniformly standardized in the literature. | (Vasudevan et al., 2024) |
| **Skin reflectance** is the wavelength-dependent fraction of incident light remitted from the skin under a defined illumination and measurement geometry. It is a physical property that arises from the absorption and scattering behavior of tissue and the distribution of biological chromophores such as melanin and hemoglobin. Reflectance serves as the optical basis for subsequent color perception and for inference of chromophores in the skin. | (Wang et al., 2017) |
| **Skin color** is a perceptual descriptor of how skin appears under specific lighting and viewing conditions. In research and clinical settings, it is operationalized by transforming measured reflectance or calibrated RGB data into standardized color spaces such as RGB and CIELAB. Skin color depends on both the underlying reflectance of the tissue and the properties of the illumination and observer. | (Weatherall and Coombs, 1992) |
| **Skin pigmentation** is a biological descriptor referring to the amount, type, and distribution of melanin in the epidermis, with smaller contributions from other pigments. Pigmentation shapes the broadband absorption profile of the skin and therefore influences its reflectance and perceived color. It is typically inferred from optical measurements that are sensitive to melanin's characteristic absorption rather than measured directly in vivo. | (Jablonski and Chaplin, 2017) |
| **Skin tone** is a cosmetic and sociocultural term describing a person's overall perceived shade or category (for example, "fair," "medium," "deep"). Context- and illumination-dependent, and only scientifically interpretable when operationalized (for example, via ITA from CIELAB). | (Jablonski, 2020; Wilder, 2010) |

## 2 Light–Tissue Interactions Relevant to Skin Measurement

Optical measurements quantify what happens to photons as they pass into or interact with a layered and structurally complex tissue. When incident light reaches the skin surface, some photons undergo specular reflection, a surface-only process perceived as shine and removable with cross-polarized photography. Most photons that contribute to skin appearance, however, enter the epidermis or dermis, undergo one or more scattering events, and re-emerge as diffuse reflectance (Figure 1 A). Diffuse reflectance carries the biologically relevant information about pigmentation, vascular absorption, and tissue structure. Photons may also undergo absorption by chromophores such as melanin and hemoglobin, or transmission, which is limited across the visible range for typical skin thicknesses. Because transmission of photons is minimal, in vivo color and spectrophotometric instruments are designed to measure remitted light, meaning the combined surface-reflected and subsurface-scattered photons returned to the detector (Wang et al., 2017; Vasudevan et al., 2024).

**Figure 1:**
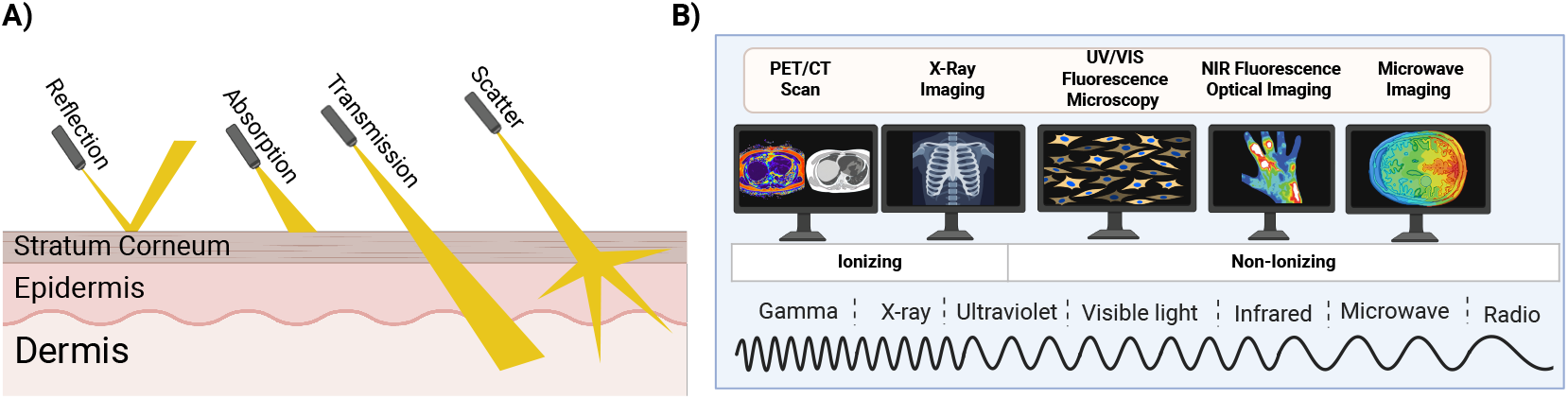
Light-skin interactions & imaging modalities across the electromagnetic spectrum. (A) Schematic representation of photon interactions with skin, showing surface reflection, subsurface scattering and absorption, and limited transmission through. Incident light interacting with skin may undergo reflection, in which photons are redirected at the tissue surface due to refractive index mismatch; scattering, in which photons are deflected multiple times within tissue by refractive index heterogeneity; or absorption, in which photon energy is taken up by tissue constituents and removed from the propagating light field. Transmission refers to photons that pass through tissue without being absorbed or back-scattered. (B) Schematic overview of imaging modalities mapped onto the electromagnetic spectrum. Modalities are positioned according to the wavelength and photon energy of the radiation they employ, from shorter-wavelength, higher-energy radiation on the left (e.g., PET/CT using gamma rays and X-rays) to longer-wavelength, lower-energy radiation on the right (e.g., infrared-based optical imaging and microwave imaging). UV/visible fluorescence microscopy and near-infrared optical imaging occupy intermediate regions of the spectrum corresponding to optical wavelengths. The spectrum is divided into ionizing and non-ionizing regimes to indicate whether photon energies are sufficient to ionize atoms in biological tissue.

In practice, these photon interactions are summarized by a single measurable quantity: the spectral reflectance of the skin. Spectral reflectance is the fraction of incident light returned from the skin at each wavelength under a specified illumination and detection arrangement (Wang et al., 2017; Seo et al., 2012; Vasudevan et al., 2024; Ly et al., 2020). Absorbance, by contrast, describes wavelength-dependent photon removal by chromophores. Because visible light does not traverse skin fully under standard in vivo measurement conditions, absorbance cannot be measured directly; instead, it is inferred from reflectance or estimated with light-transport models that incorporate the absorption spectra and spatial distributions of melanin and hemoglobin. For visible-range tools, spectral reflectance is therefore the operational signal from which chromophore-related information must be interpreted.

Photons at different wavelengths carry different energies, and these energy differences produce different patterns of interaction with tissue. This is the basis for the range of imaging and therapeutic modalities across the electromagnetic spectrum (Figure 2B). At the high-energy end are X-rays and gamma rays, which are classified as ionizing radiation because they can remove electrons from atoms and create ionized molecules. These wavelengths support imaging modalities such as radiography, computed tomography (CT), and nuclear medicine techniques including positron emission tomography (PET) and single-photon emission computed tomography (SPECT). At lower energies are the ultraviolet (UV), visible, and near-infrared (NIR) regions. UV spans both non-ionizing and ionizing wavelengths: the shortest-wavelength UV (UVC and part of UVB) can ionize molecules, while longer-wavelength UV interacts with tissue primarily through absorption and scattering. Across the visible and NIR ranges, photon interactions with skin are likewise governed mainly by absorption and scattering rather than ionization. These wavelength regions support a broad range of optical approaches, including UV photography, clinical photography, and visible-or NIR-range spectrophotometry. They are also used therapeutically because different wavelengths interact preferentially with different tissue components. For example, visible wavelengths interact strongly with hemoglobin and can therefore target superficial vascular structures, whereas NIR wavelengths interact more weakly with melanin and penetrate more deeply, and mid-infrared wavelengths are absorbed strongly by water and form the basis of ablative and fractional resurfacing systems (Anderson and Parrish, 1983; Alexiades-Armenakas et al., 2008; Gan and Graber, 2013). At the longest infrared wavelengths (far-IR), imaging relies primarily on emitted thermal radiation, which can be used to assess surface temperature patterns related to inflammation or perfusion (Lahiri et al., 2012).

**Figure 2:**
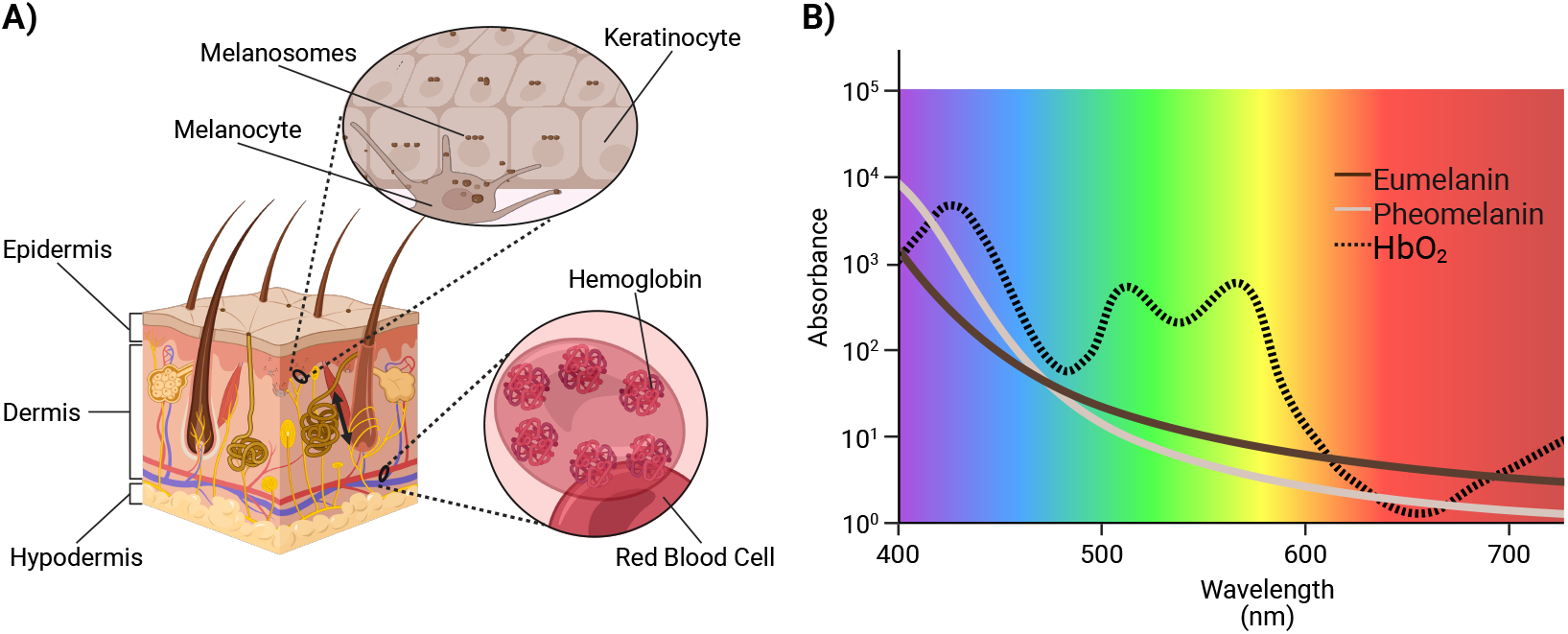
General location and absorption of major chromophores in skin. (A) A skin cross-sectional diagram il-lustrating the cellular sources and spatial distribution of key chromophores. Melanin is produced by melanocytes in the basal epidermis and packaged into melanosomes that are transferred to keratinocytes, while hemoglobin is contained within red blood cells in the dermal vasculature.(B) Absorption spectra of eumelanin, pheomelanin, and hemoglobin across the visible wavelength range. Eumelanin exhibits strong, broadband absorption that decreases monotonically with increasing wavelength, whereas pheomelanin shows weaker, more spectrally structured absorption, particularly at shorter visible wavelengths. Oxy- and deoxyhemoglobin display distinct, wavelength-specific absorption features. Spectral overlap between melanin subtypes and hemoglobin underlies challenges in disentangling pigmentation and erythema signals from diffuse reflectance measurements.

## 3 Chromophores in Skin and Their Spectral Behavior

Human skin color arises from the biological distribution, composition, and organization of pigments within the epidermis and dermis. Variation in the packaging of these chromophores define the true phenotypic variation underlying skin appearance, independent of how that appearance is ultimately perceived, categorized, or measured. To translate these biological differences into the optical domain, however, we also require quantitative descriptors of how much of each chromophore is present. Because chromophores occupy physical space within defined anatomical compartments, one universal way to express their abundance is as the volume fraction of a given layer taken up by that pigment. Volume fraction provides a common quantitative language that links histologic measurements, biochemical assays, and optical models: it describes the proportion of epidermis occupied by melanin-containing melanosomes, or the proportion of superficial dermis occupied by intravascular blood, regardless of measurement modality (see Table 2).

**Table 2:** Representative chromophore volume fractions in human skin. Values for melanin and dermal blood are derived from optical measurements combined with light-transport modeling; eumelanin and pheomelanin composition is based on chemical analysis of epidermal tissue samples.

| Chromophore | Volume fraction range | Basis | Key references |
| --- | --- | --- | --- |
| Total melanin | 5–20% on forearm (Fitz I–VI); higher values reported for very darkly pigmented skin at other sites. | Saager et al. estimate 5–20% melanin volume fraction across Fitzpatrick I–VI on the top and inner forearm by fitting optical models to in vivo multiphoton microscopy and spatial frequency-domain spectroscopy measurements. | (Saager et al., 2015) |
| Eumelanin | Majority of epidermal melanin in all skin; approximately 92–98% of total melanin by mass. | Alaluf et al. quantify eumelanin and pheomelanin in epidermal samples using chemical degradation and biochemical assays; Thody et al. detect both pigment types in all samples using similar methods. | (Alaluf et al., 2002; Thody et al., 1991) |
| Pheomelanin | Present in all skin; approximately 2–8% of total melanin by mass in darkly pigmented skin. | Thody et al. detect pheomelanin in all epidermal samples using biochemical assays; Alaluf et al. report pheomelanin as a minor component across all groups using chemical degradation methods, similarly to Del Bino et al. | (Thody et al., 1991; Alaluf et al., 2002; Del Bino et al., 2015) |
| Dermal blood (rest) | Whole dermis: 0.2–0.6% typical (0.08–1.2% measured); papillary: 0.9–4%; reticular: 0.7–1.9%. | Fredriksson et al. estimate whole-dermis blood by fitting optical models to laser Doppler flowmetry signals (0.08% forearm, 1.2% fingertip at rest). Verdel et al. estimate layer-specific blood content by fitting optical models to combined photothermal and reflectance measurements. | (Verdel et al., 2019; Fredriksson et al., 2008) |

Melanin is the principal absorber in the epidermis and the primary determinant of broadband visible absorption. Human epidermal melanin consists of two pigments generated through mixed melanogenesis: eumelanin, which predominates across skin phototypes, and pheomelanin, which is present in smaller but measurable amounts (Ito and Wakamatsu, 2003).

Notably, while the pheomelanin fraction in hair is much higher in fair and red-haired individuals (Vincensi et al., 1998; Wakamatsu et al., 2002) this does not translate to a qualitatively different epidermal reflectance: even in these phenotypes, visible skin color is dominated by eumelanin, with elevated pheomelanin manifesting primarily as increased UV sensitivity rather than altered skin appearance (Hennessy et al., 2005).

These biochemical distinctions translate into characteristic spectral behavior (Figure 2B). Eumelanin absorbs strongly at short wavelengths, with absorption decreasing smoothly toward the red end of the visible spectrum, producing the broadband attenuation that darkens skin and suppresses blue-green reflectance. Pheomelanin absorbs less strongly overall and has a more yellow-red spectral profile. Given its small fraction in epidermis (Table 2), its direct contribution to visible reflectance is minor; it becomes optically relevant mainly when total melanin is low, and more importantly through its influence on UV response rather than steady-state skin color.

Hemoglobin in the papillary and upper reticular dermis is the major source of narrow-band absorption in the visible spectrum (Figure 2A). Because hemoglobin is confined to the superficial cutaneous microvasculature, its optical contribution depends on vessel density, blood volume, and oxygenation state, which vary across anatomical sites and physiological conditions. Oxygenated hemoglobin exhibits a characteristic twin-peaked absorption profile near 542 nm and 578 nm (Figure 2B), while deoxygenated hemoglobin shows a single broader peak near 555 nm (Zijlstra et al., 2021). These oxygenation-dependent spectral differences underlie clinical perceptions of redness and enable optical assessment of tissue oxygen saturation (Lister et al., 2012; Meglinski and Matcher, 2002). The spectral overlap between melanin and hemoglobin absorption, particularly at shorter visible wavelengths, underlies one of the central challenges in skin optics: disentangling pigmentation and erythema signals from diffuse reflectance measurements.

Other chromophores contribute to visible-range reflectance but exert smaller effects under typical conditions. Carotenoids absorb primarily in the blue-green region and can shift skin appearance toward yellow in specific dietary or pathophysiological contexts. Water and lipids have minimal absorption in the visible range but dominate specific near-infrared wavelengths, where extended-range imaging becomes sensitive to hydration and deeper dermal structure. Under broad-spectrum visible illumination of healthy skin, these pigments play a secondary role relative to melanin and hemoglobin (Lister et al., 2012).

Taken together, these chromophores produce the characteristic spectral signature of skin: melanin shapes the broadband baseline, hemoglobin adds wavelength-specific vascular features, and minor pigments contribute smaller context-dependent effects. These combined interactions define the visible-range reflectance patterns that underlie both clinical appearance and optical measurement.

## 4 Tools for Measuring Skin Color and Their Reported Outputs

All instruments used to quantify skin color operate on the same principle: illuminate the skin with a controlled light source and measure the light that returns to a detector. What distinguishes devices is how they sample that returning light. Figure 3 compares the major instrument classes by spatial information, spectral capability, output type, and cost.

**Figure 3:**
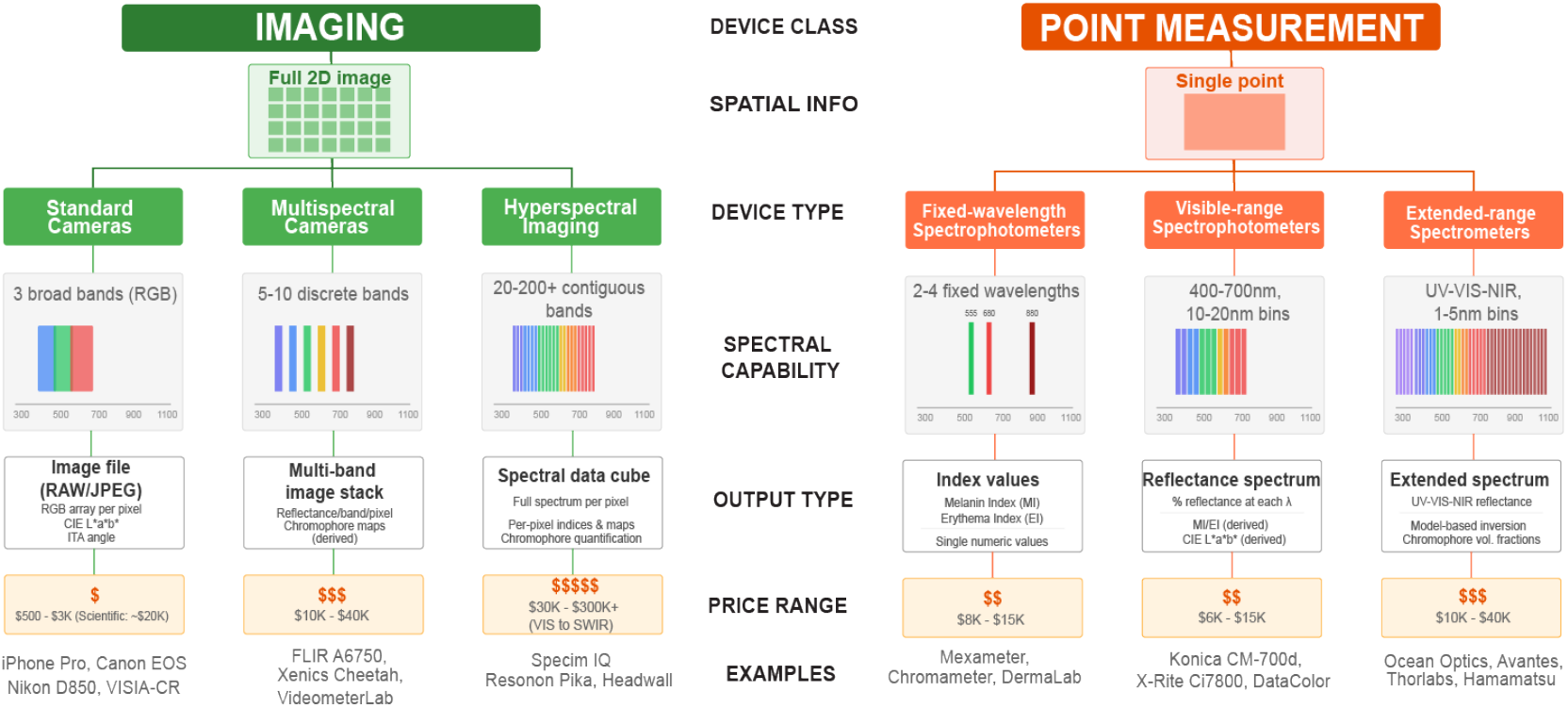
Tools for measuring skin color. An illustrative comparison of commonly used skin-measurement technologies, organized by device class, spectral capability, output type, and approximate cost range. Imaging-based systems, including multispectral and hyperspectral cameras, acquire spatially resolved reflectance data across multiple wavelengths, enabling chromophore mapping and spectral analysis at the pixel level. In contrast, fixed-wavelength spectrophotometric devices provide point measurements and output index-based summaries (e.g., melanin or erythema indices). Differences in spectral resolution, spatial information, and analytical complexity across device classes have important implications for measurement bias, accessibility, and interpretability.

The primary distinction is between imaging devices, which capture a full 2D field of view, and point-based devices, which measure reflectance from a single small area of skin. Within each category, instruments vary in how finely they resolve the spectrum: from three broad RGB channels to multispectral systems with 5-10 discrete bands to hyperspectral imaging with hundreds of contiguous wavelength bands, and from a few fixed wavelengths to continuous visible and extended-range coverage for point-based devices. As spectral resolution increases, so does the potential to separate chromophore contributions, but so does the cost of the instrument and analytical complexity of the data.

The choice between imaging and point measurement depends on the question being addressed. Imaging systems provide spatial context essential for documenting lesion extent or comparing affected to unaffected regions. Point-based devices are well suited to longitudinal tracking or standardized site comparisons where a single representative measurement suffices.

### Output Metrics and Their Interpretation

The numerical values reported by these instruments take several forms. RGB values are intuitive but device-dependent and conflate contributions from all chromophores. CIELAB provides a perceptually uniform color space useful for quantifying color change, but still describes appearance rather than biology. Individual Typology Angle (ITA) reduces skin color to a single number at the cost of discarding information about redness entirely, making it insensitive to erythema (Chardon et al., 1991; Ly et al., 2020).

Melanin Index and Erythema Index are the most commonly reported outputs from fixed-wavelength spectrophotometers. The Melanin Index is typically derived from absorbance at red or near-infrared wavelengths where hemoglobin absorption is minimal; the Erythema Index from absorbance in the green region where hemoglobin absorbs strongly. These indices are reproducible within a device and useful for tracking relative changes, but they are not standardized across manufacturers and, critically, they are not independent measurements of chromophore concentration. Melanin absorbs at all visible wavelengths including those used for erythema estimation, and hemoglobin absorption extends into the red region used for melanin estimation.

Raw reflectance spectra preserve the full information content of the measurement. Because all other metrics are mathematically derived from spectral reflectance, raw spectra can be transformed post hoc into CIELAB, ITA, or Melanin/Erythema Indices (Vasudevan et al., 2024; Seo et al., 2012; Wang et al., 2017). The trade-off is that spectral data are higher-dimensional and require explicit calibration and modeling.

From this perspective, all commonly used metrics represent deliberate reductions of a higher-dimensional signal into coordinates intended to capture the most salient variation (Clarys et al., 2000; Shriver and Parra, 2000; Takiwaki et al., 2002; Weatherall and Coombs, 1992). Whether these dimensionality reduction approaches reflect biologically salient dimensions of variation remains an important question. Direct validation of optical melanin or erythema measures against tissue-level endpoints has been reported in only a limited number of studies, with most work relying on subjective or semi-quantitative clinical reference standards (Vasudevan et al., 2024; Weir et al., 2024). Understanding what these skin reflectance metrics successfully capture and what they lose requires examining the physics of light-tissue interaction.

## 5 Fundamental Limits of Visible-Range Skin Measurement

Interpreting skin reflectance requires understanding what happens when light encounters multiple absorbing substances simultaneously. The Beer-Lambert law describes this interaction: absorbance at any wavelength is proportional to the absorber’s concentration and the path length light travels through the medium. When multiple chromophores are present, their absorbances sum. This additivity seems straightforward, but it has a critical implication for skin measurement: the practical question is not whether a chromophore contributes to the spectrum, but whether its contribution can be distinguished from other contributions and from measurement noise.

Figure 4 illustrates two scenarios that produce very different outcomes despite both following Beer-Lambert additivity. In Panel A, two chromophores with non-overlapping absorption peaks are mixed in increasing concentrations. A blue solution absorbs in the red/orange region of the spectrum; a red dye absorbs in the blue/green region. As the blue dye is added, the solution becomes purple, and critically, both absorption peaks remain visible in the spectrum. Because the peaks occupy different wavelength ranges, each can be measured independently. The chromophores are spectrally separable, and increasing the concentration of one does not obscure information about the other.

**Figure 4:**
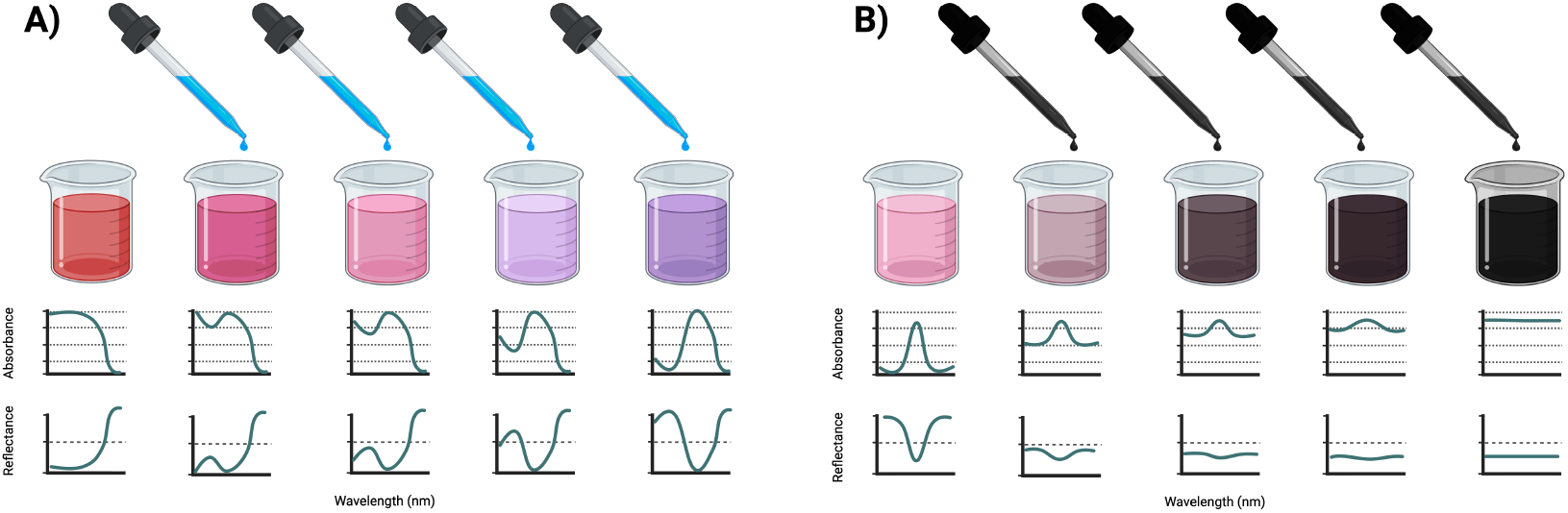
Schematic representation of Beer-Lambert additivity. A conceptual demonstration of how spectral overlap influences chromophore detectability. (A) Non-overlapping absorbers remain independently measurable with increasing concentration. (B) A broadband absorber elevates baseline absorbance and suppresses the relative prominence of narrow spectral features, limiting detectability despite additive absorption.

Panel B illustrates a different situation. Here, a chromophore with a narrow absorption peak (a pink solution absorbing in the green region) is mixed with a broadband absorber (black ink, which absorbs across all visible wavelengths). As black ink is added, total absorbance rises at every wavelength. The narrow absorption peak remains present in the mathematical sense, contributing the same absolute absorbance as before. However, that fixed contribution becomes an increasingly small fraction of the total signal. The contrast between the peak wavelength and surrounding wavelengths, which is what encodes the presence of the narrowband chromophore, shrinks until it falls below the detection threshold of the measurement system. The solution appears uniformly black, and no amount of careful measurement can recover information about the pink chromophore’s presence.

This second scenario is directly analogous to measuring hemoglobin in skin with varying melanin content. Hemoglobin produces characteristic absorption peaks in the 540-580 nm range. Melanin absorbs broadly across the visible spectrum, with absorption increasing toward shorter wavelengths. As melanin concentration increases, it raises the baseline absorbance at all wavelengths, progressively reducing the relative prominence of hemoglobin’s spectral features. The hemoglobin is still present and still absorbing light, but its signature becomes undetectable against the elevated baseline.

The situation in skin is further complicated by the layered distribution of these chromophores. Melanin is concentrated in the epidermis, while hemoglobin circulates in the dermal vasculature below. Light must pass through the melanin-containing epidermis to reach hemoglobin, then traverse it again on the return path to the detector. This double-pass geometry means that hemoglobin’s signal is attenuated by melanin twice, causing the signal to decrease faster than a simple additive model would predict. The attenuation is also wavelength-dependent: shorter wavelengths, where melanin absorption is strongest, are most affected.

These physical constraints have direct consequences for the indices commonly used in skin measurement. The Melanin Index and Erythema Index are calculated from absorbance values at wavelengths chosen to emphasize melanin or hemoglobin contributions, but both wavelength ranges receive contributions from both chromophores. In lightly pigmented skin, hemoglobin features are prominent and the Erythema Index primarily reflects vascular changes. As pigmentation increases, melanin’s contribution at erythema wavelengths grows while hemoglobin’s relative contribution shrinks. At high melanin concentrations, the Erythema Index may be dominated by melanin absorption, with hemoglobin features falling below the noise floor entirely. This is not a calibration problem that better instruments could solve. It is a physical consequence of overlapping absorption spectra combined with the layered anatomy of skin.

## 6 Empirical Case Study: Broadband Reflectance Demonstration of Optical Limits

The recently published International Skin Spectra Archive (ISSA) provides a unique opportunity to evaluate the effect of pigmentation and structural variation in skin anatomy, as it contains over 15,000 visible-range spectra taken across body sites from more than 2,000 participants across multiple countries (Figure 5A; Lu et al. 2025a). Using this dataset, we demonstrate how the physical principles described above manifest in empirical data by focusing on the distribution of pigmentation, defined by Melanin Index (MI), measured on participant faces and palms (Figure 5B). The number of individuals with forehead and or cheek readings was 2,101, while palm reflectance spectra were available for 776 individuals.

**Figure 5:**
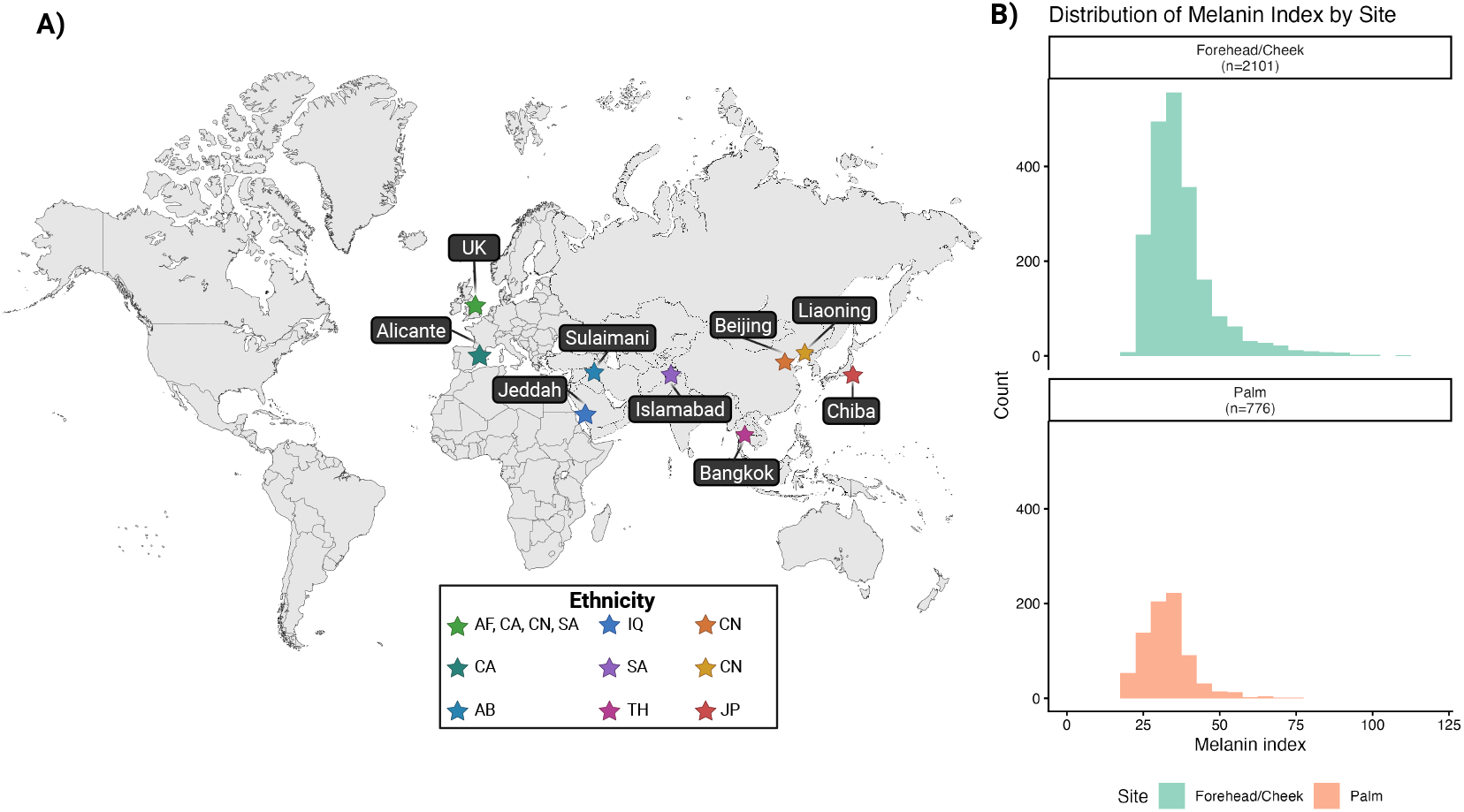
International Skin Spectra Archive (ISSA) participant map and melanin index distribution. (A) Map showing ISSA participant locations and reported ethnicities, including African (AF), Caucasian (CA), Chinese (CN), and South Asian (SA) individuals from four U.K. cities (Sheffield, Manchester, Leeds, Liverpool); CA from Alicante, Spain; Arab (AB) from Jeddah, Saudi Arabia; Iraqi (IQ) from Sulaimani, Iraq; SA from Islamabad, Pakistan; Thai (TH) from Bangkok, Thailand; CN from Beijing and Liaoning, China; and Japanese (JP) from Chiba, Japan. (B) Histograms showing the distribution of melanin index values for forehead or cheek (top) and palm (bottom). Original data published in *Scientific Data* and available via figshare (Lu et al., 2025b).

To facilitate visual comparison of spectral features across the pigmentation range between individuals, we take facial MI as a representative value of an individual’s level of pigmentation and classify individuals according to the Eumelanin Human Skin Color Scale (EHSC) (Dadzie et al., 2022): Low (MI < 25), Intermediate Low (MI 25 to 50), Intermediate (MI 50 to 75), Intermediate High (MI 75 to 100), and High (MI > 100). This quantitative classification allows us to examine how spectral structure changes systematically across the gradient of skin pigmentation. Looking at raw reflectance values, we see that the spectra appear progressively flatter as skin becomes more pigmented (Figure 6A). The characteristic W-shape around 555nm, attributed to the signal of hemoglobin and key to the calculation of Erythema Index (EI) with spectrophotometers, becomes less prominent as skin pigmentation increases.

**Figure 6:**
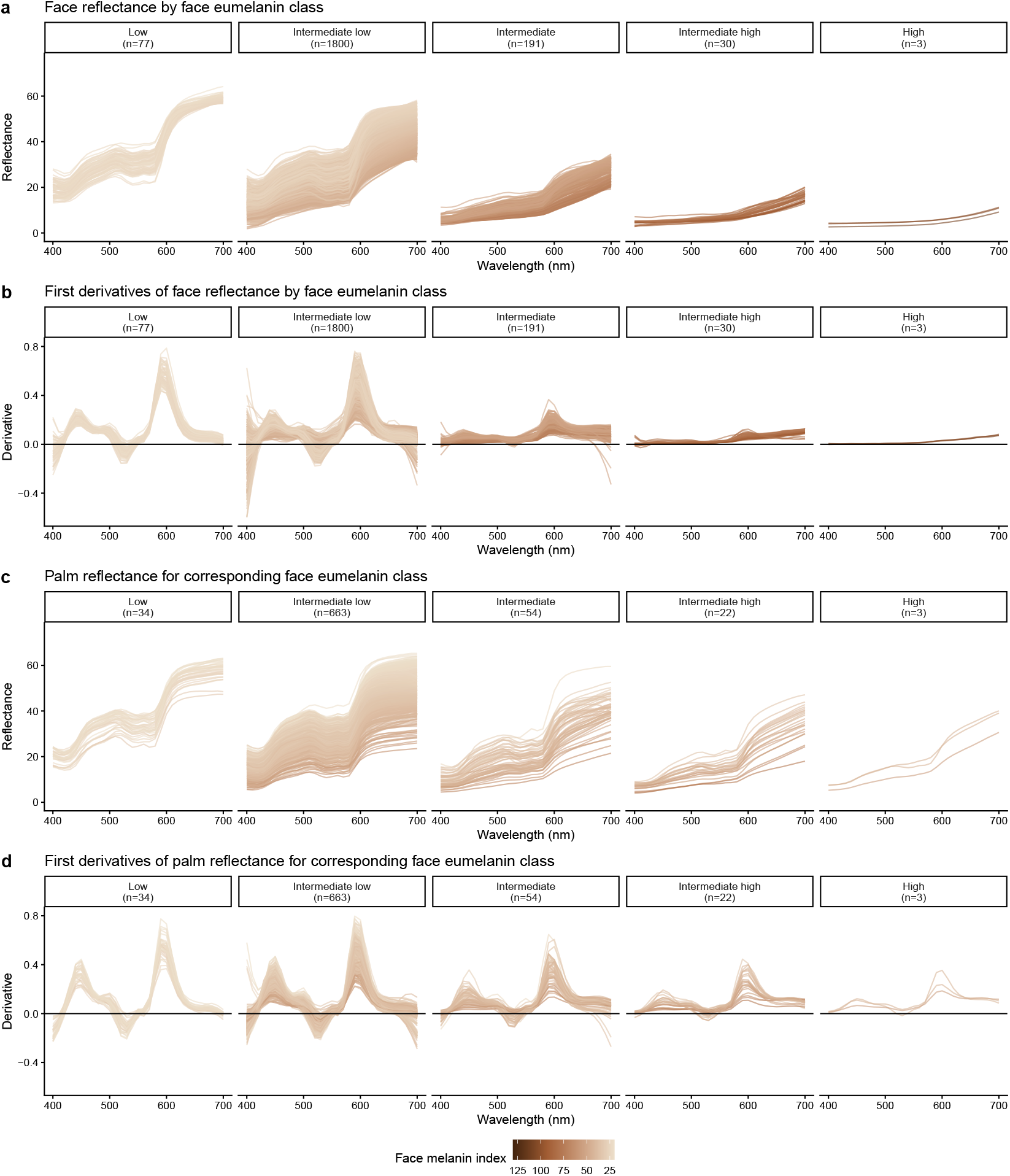
Facial and palm reflectance spectra and first derivatives across facial eumelanin classes. Raw reflectance spectra measured on the face (forehead or cheek) are shown in (A), and corresponding palm spectra from the same individuals are shown in (B), with individuals grouped by facial eumelanin class defined using Melanin Index (MI). First derivatives of the reflectance spectra are shown for face in (C) and palm in (D). The first derivative represents the change in reflectance with wavelength and was calculated from each spectrum using local polynomial regression. Spectra with pronounced hemoglobin-related structure yield derivatives with large peak and valley amplitudes, whereas flatter spectra yield derivatives closer to zero. As pigmentation increases, hemoglobin-related derivative features between 500 and 650 nm are strongly attenuated in facial spectra, while corresponding palm spectra retain greater derivative structure across the same pigmentation classes.

To better characterize the spectral shape, we calculated the first derivative of each spectral curve (Figure 6B). A spectrum with prominent peaks and valleys will have a derivative with large amplitude, whereas a flat spectrum will have a derivative near zero. What becomes apparent in this representation is that the distinct signal from hemoglobin, namely the valleys and peaks of the derivative values between 500 and 650 nm, is severely attenuated in facial spectra as pigmentation increases (Figure 6B). Interestingly, the corresponding palm spectra and derivatives for individuals in those same EHSC classes show less attenuation of these features (Figure 6C and 6D). This pattern could partially be explained by the fact that an individual’s palms and soles are generally lighter than the rest of their skin due to the anatomy of glabrous skin.

As a more formal means of examining whether this difference reflects pigmentation alone or an additional site-specific effect, we quantified hemoglobin-related signal amplitude in two ways: the difference between the peak and valley of the first-derivative spectrum within 500 to 650 nm, and the reflectance difference between 680 nm and approximately 555 nm, the latter forming the basis of EI. For both measures, signal amplitude declined with increasing MI at both anatomical sites, following a power-law relationship (Figure 7).

**Figure 7:**
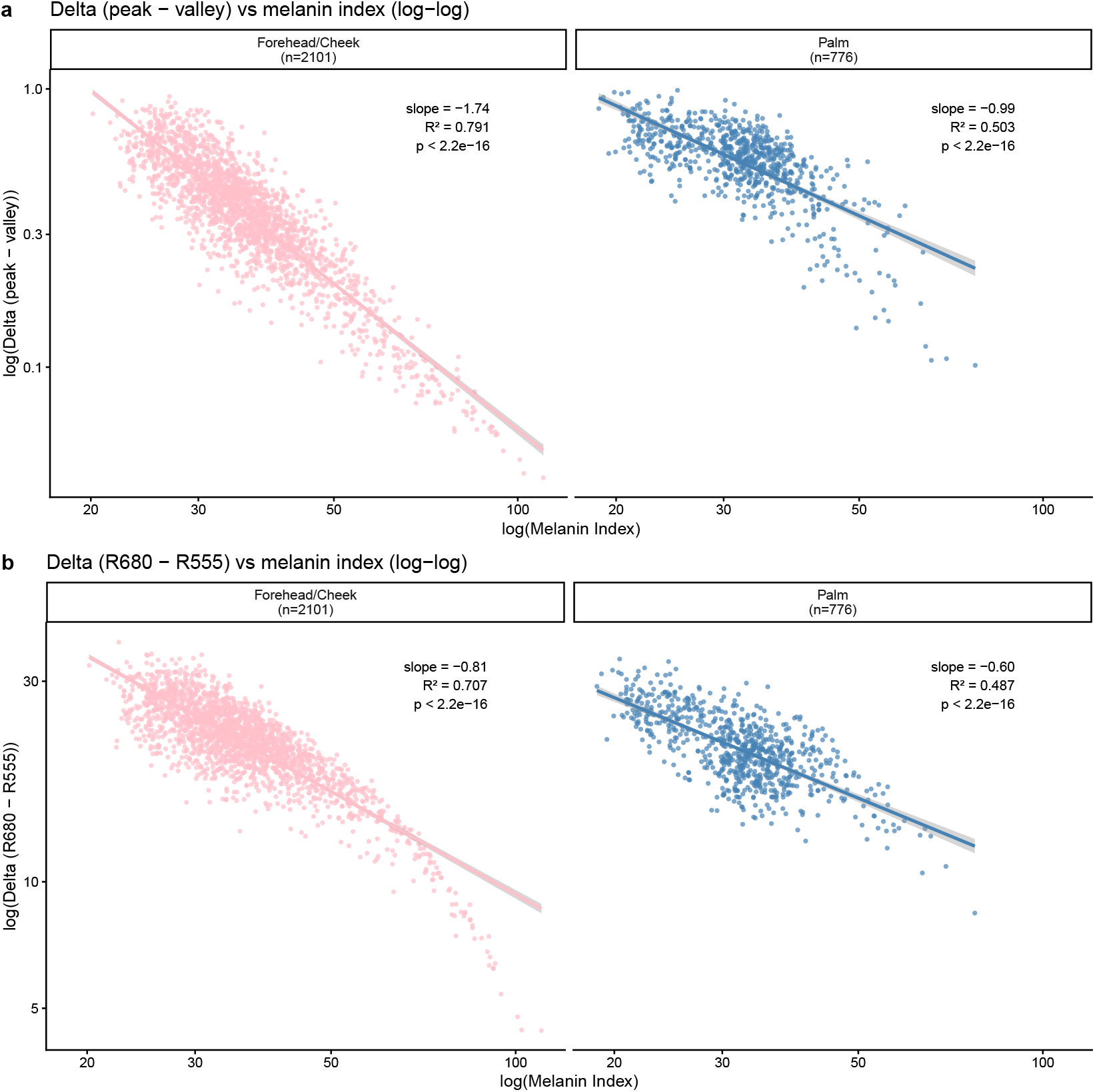
Pigmentation-dependent attenuation of hemoglobin-related signal differs by anatomical site. (A) Log–log relationship between Melanin Index (MI) and peak–valley amplitude of the first-derivative spectrum (500 to 650 nm) for face (forehead or cheek) and palm. (B) Log–log relationship between MI and reflectance difference between 680 nm and approximately 555 nm (R680–R555), which is the basis for the Erythema Index. Points represent individual measurements and lines show fitted relationships within each site. In both cases, signal amplitude declines with increasing pigmentation, but attenuation is steeper on the face than on the palm. Differences in scaling by anatomical site were formally tested using linear mixed-effects models with a random intercept for subject, comparing models with and without site-specific slopes.

Importantly, however, the rate of attenuation differed by anatomical site even after accounting for pigmentation. Mixed-effects models accounting for repeated measurements within individuals showed that anatomical site significantly modified the relationship between MI and hemoglobin-related signal amplitude for both peak to valley derivative delta and reflectance delta, compared to models including MI alone (likelihood ratio tests, *p* < 2.2 × 10^−16^ for both outcomes). In practical terms, hemoglobin-related contrast diminishes more rapidly with increasing pigmentation on the face than on the palm, indicating that the observed difference cannot be explained solely by the lighter pigmentation of glabrous skin. The complete statistical analyses can be found on https://lasisilab.github.io/SkinOptics/JID_SkinOptics.html.

This attenuation has particular clinical significance because of what each chromophore represents physiologically. Melanin content is relatively stable within an individual over time. Hemoglobin-related signals, by contrast, are dynamic, reflecting changes in blood flow, oxygenation, and inflammatory state in response to physiological perturbation, disease processes, and treatment. The erythema index is intended to track these changes. As pigmentation increases, however, this dynamic signal is increasingly masked by the stable melanin background, reducing the effective sensitivity of erythema-based measurements. The result is a progressive loss of contrast that limits the ability to detect clinically meaningful change in more highly pigmented facial skin.

## 7 Moving Forward: Improving Skin Color Measurement and Interpretation

What the visible spectrum can and cannot tell us about skin is not a question of instrument quality or image resolution. It is a question of physics. The same optical entanglement that limits spectrophotometric indices also limits what clinical photography can capture and what AI systems can learn from photographic data. When we build tools without understanding these limits, we do not overcome them—we encode them.

Visible-range reflectance measurements capture real optical information, and the shift toward objective, quantitative assessment represents genuine progress. But interpreting that information requires understanding the physical constraints imposed by chromophore absorption and tissue structure. Melanin Index offers a practical framework for navigating these constraints, grounded in the optical property that actually determines measurement performance.

When working with existing datasets, MI distributions can reveal sampling gaps that demographic summaries obscure. Traditional markers of sample diversity do not guarantee adequate coverage of the pigmentation range that matters for optical measurement. A sample can appear diverse by demographic criteria while remaining sparse at the high end of the MI distribution, as we observed even in a large multi-country dataset. Yet this is precisely where optical entanglement is most severe, where spectral features are most attenuated, and where current indices are least validated. When the high-MI range is underrepresented, conclusions should be appropriately qualified. Where full spectra are available, derivative or other shape-based analyses can reveal hemoglobin-related structure that persists in darker skin even when standard indices show no signal.

When collecting new data, sampling should explicitly target balanced representation across the MI distribution, with particular attention to individuals with MI above 75, where optical entanglement is most apparent and existing data are scarcest. Experimental designs that incorporate controlled physiological perturbation, multi-site sampling within individuals, and consistent reporting of measurement conditions will provide the empirical foundation needed to validate approaches in darker skin. At minimum, publications should report the MI distribution of participants, anatomical sites measured, and device specifications.

Some applications will require capabilities beyond visible-range reflectance altogether. Extending into the ultraviolet offers improved melanin characterization. Extending into the near-infrared and short-wave infrared offers access to deeper tissue structures and perfusion measurements less attenuated by melanin. Biophysical modeling can guide algorithm development, but its value depends on parameter ranges validated for darker skin and, ultimately, on correlating optical outputs with histological ground truth.

Collectively, these considerations underscore the need to translate optical objectivity into measurements that are biologically informative, interpretable, and valid across the full range of human skin pigmentation.

## Acknowledgements

We thank DerMEDit (www.dermedit.com) for critical review and discussions related to the development of this manuscript.

## CRediT Contribution Statement

Concepualization: TL

Formal Analysis: TL, JH

Supervision: TL, JK

Visualization: TL, YP

Writing-original draft: TL, YP

Writing-review & editing: TL, YP, JH, JK, BJ

